# WWOX-Mediated Degradation of AMOTp130 Negatively Affects Egress of Filovirus VP40 VLPs

**DOI:** 10.1101/2021.11.19.469355

**Authors:** Jingjing Liang, Gordon Ruthel, Bruce D. Freedman, Ronald N. Harty

**Author notes:** <u>Corresponding Author: Dr. Ronald N. Harty</u>, <u>Professor</u>, Department of Pathobiology, School of Veterinary Medicine, University of Pennsylvania, 3800 Spruce Street, Philadelphia, PA 19104, USA.

## Abstract

Ebola (EBOV) and Marburg (MARV) viruses continue to emerge and cause severe hemorrhagic disease in humans. A comprehensive understanding of the filovirus-host interplay will be crucial for identifying and developing antiviral strategies. The filoviral VP40 matrix protein drives virion assembly and egress, in part by recruiting specific WW-domain-containing host interactors via its conserved PPxY Late (L) domain motif to positively regulate virus egress and spread. In contrast to these positive regulators of virus budding, a growing list of WW-domain-containing interactors that negatively regulate virus egress and spread have been identified, including BAG3, YAP/TAZ and WWOX. In addition to host WW-domain regulators of virus budding, host PPxY-containing proteins also contribute to regulating this late stage of filovirus replication. For example, angiomotin (AMOT) is a multi-PPxY-containing host protein that functionally interacts with many of the same WW-domain-containing proteins that regulate virus egress and spread. In this report, we demonstrate that host WWOX, which negatively regulates egress of VP40 VLPs and recombinant VSV-M40 virus, interacts with and suppresses the expression of AMOT. We found that WWOX disrupts AMOT’s scaffold-like tubular distribution and reduces AMOT localization at the plasma membrane via lysosomal degradation. In sum, our findings reveal an indirect and novel mechanism by which modular PPxY/WW-domain interactions between AMOT and WWOX regulate PPxY-mediated egress of filovirus VP40 VLPs. A better understanding of this modular network and competitive nature of protein-protein interactions will help to identify new antiviral targets and therapeutic strategies.

**IMPORTANCE:** Filoviruses (Ebola [EBOV] and Marburg [MARV]) are zoonotic, emerging pathogens that cause outbreaks of severe hemorrhagic fever in humans. A fundamental understanding of the virus-host interface is critical for understanding the biology of these viruses and for developing future strategies for therapeutic intervention. Here, we reveal a novel mechanism by which host proteins WWOX and AMOTp130 interact with each other and with the EBOV matrix protein VP40 to regulate VP40-mediated egress of virus like particles (VLPs). Our results highlight the biological impact of competitive interplay of modular virus-host interactions on both the virus lifecycle and the host cell.

## INTRODUCTION

Ebola (EBOV) and Marburg (MARV) viruses are emerging pathogens that can cause severe hemorrhagic disease in humans, and these emerging pathogens remain a global public health threat that warrant urgent development of antiviral therapeutics (1–3). Toward this end, a more in-depth understanding of the interplay between the host and EBOV/MARV proteins would be beneficial both for our understanding of the fundamental molecular mechanisms of the virus lifecycle, and for identifying new targets and strategies for antiviral development.

Our focus is on the filovirus VP40 protein, which is the major structural protein that drives virion assembly and egress. Expression of VP40 alone is sufficient for the formation and egress of virus-like particles (VLPs) that mimic the morphology of authentic virus particles (4–10). In our investigations of the VP40/host interactome, we identified a series of specific host interactors that either positively or negatively regulated the budding process (11–17). These host proteins contain one or more modular WW-domains that interacted with the conserved PPxY Late (L) domain motif at the N-termini of both EBOV VP40 (eVP40) and MARV VP40 (mVP40). The PPxY L-domain plays a key role in promoting efficient release of VLPs and live virus, in part, by hijacking or recruiting host factors that enhance VLP or virus egress from the plasma membrane (18). For example, the VP40 PPxY motif interacts with several members of the HECT family of E3 ubiquitin ligases, such as Nedd4, Itch, WWP1, and Smurf2 which facilitate mono-ubiquitination of VP40 and subsequent engagement and re-localization of the host ESCRT machinery to the site of virus budding at the plasma membrane to enhance virus-cell separation and release of VLPs or infectious virus (8, 11, 16, 17). In contrast to these positive regulators of virus budding, we have identified more recently a growing list of novel WW-domain interactors that negatively regulate egress, such as host proteins BAG3, YAP/TAZ, and WWOX (12, 13, 15). Notably, these negative regulators of budding are multifunctional proteins that regulate diverse pathways/processes in the cell including, apoptosis/autophagy, transcription, cytoskeletal dynamics, cell migration/morphology, and tight junction formation (19–24). Moreover, regulation of these diverse pathways by WW-domain containing BAG3, YAP/TAZ, and WWOX is achieved, in part, via their interactions with host PPxY-containing proteins, such as angiomotin (AMOT) (25–30). Notably, AMOT contains three N-terminal PPxY motifs, two of which are identical to that in eVP40. Thus, we speculated that the modular mimicry of the viral and host PPxY motifs could lead to competitive interactions with host WW-domain containing proteins resulting in meaningful biological consequences for both the virus and the host. Indeed, we recently demonstrated that expression of AMOT had a positive influence on egress of both VP40 VLPs and live EBOV and MARV by counteracting the negative effects of YAP and WWOX (12, 13, 31).

In this report, we investigated further the interplay among AMOT, WWOX, and eVP40/mVP40 to understand the molecular mechanism by which these host proteins and modular interactions regulate budding. Our results revealed that WWOX interacts with and regulates expression and intracellular distribution of AMOT, and that this interplay between AMOT and WWOX contributes mechanistically to the efficiency of VP40 VLP egress. Specifically, we found that expression of WWOX reduced the levels of AMOT in the cytoplasm and at the plasma membrane, as well as modified the intracellular localization of AMOT, and that these changes in AMOT expression and localization correlated with a decrease in VP40 VLP egress. Moreover, expression of WWOX led to lysosomal degradation of AMOT and a concomitant decrease in VP40 VLP egress. In sum, these data highlight a novel, complex network of modular PPxY/WW-domain interactions that may impact both the host and the late stages of filovirus egress and spread.

## RESULTS

### WWOX interacts with and reduces AMOTp130 expression levels

We demonstrated previously that expression of full-length, endogenous AMOT (AMOTp130) was important for the efficient egress of both VP40 VLPs and for live infectious EBOV and MARV, whereas expression of WWOX inhibited egress of VP40 VLPs from HEK293T cells. In light of the contrasting roles for PPxY-containing AMOTp130 and WW-domain containing WWOX in regulating VP40-mediated egress, we first sought to determine whether WWOX and AMOTp130 interact. Toward this end, we transfected HEK293T cells with expression plasmids for WWOX (myc-tagged) and AMOTp130 (HA-tagged) and used an IP/Western approach to demonstrate a strong interaction between the two exogenously expressed proteins (Fig. 1A). In addition, we used the same approach to demonstrate that exogenously expressed WWOX interacted with endogenous AMOTp130 (Fig. 1B).

**FIG. 1.**
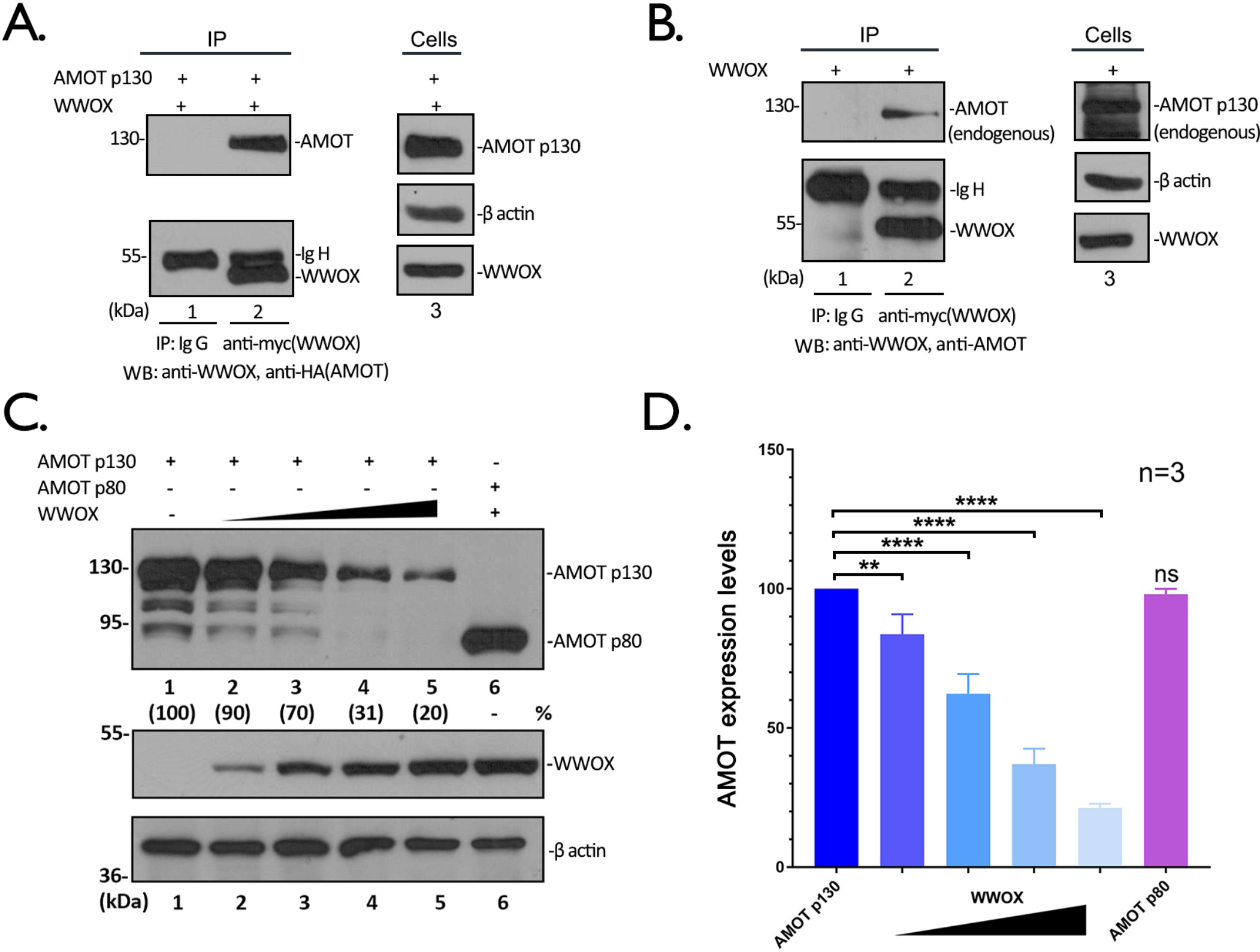
WWOX interacts with AMOT and reduces its expression levels in a dose-dependent manner. **A)** Extracts from HEK 293T cells transfected with myc-tagged WWOX and HA-tagged AMOT were immunoprecipitated (IP) with either normal mouse IgG or anti-myc antibody. AMOT and WWOX were detected in the precipitates by Western blot (WB). Expression controls for AMOT, WWOX and β-actin are shown at the left panel. **B)** Extracts from HEK 293T cells transfected with WWOX alone were IP with either normal mouse IgG or anti-myc antibody. Endogenous AMOT and WWOX were detected in the precipitates by WB. **C)** HEK293T cells were transfected with AMOTp130 or p80 plus vector (-) or increasing amounts of WWOX. The indicated proteins were detected by WB and AMOT expression levels were quantified () using NIH Image-J. The amounts of AMOT in control cells (lane 1) were set at 100%. The relative levels of AMOT are shown in (). **D)** Quantification of the AMOT levels from three independent experiments. Statistical significance was analyzed by a one-way ANOVA. ns: not significant, **= p<0.01, ****= p<0.0001.

Next, we asked whether transfecting HEK293T cells with increasing amounts of WWOX plasmid would affect expression levels of AMOTp130 (Fig. 1C). Intriguingly, we observed a dose-dependent decrease in the levels of AMOTp130 as the amount of transfected WWOX plasmid was increased (Figs. 1C and 1D). Notably, WWOX expression did not reduce the level of exogenously expressed AMOTp80; an N-terminally deleted isoform of AMOTp130 that lacks all of the PPxY motifs. Together, these data show that WWOX interacts with AMOTp130 and that increased expression of WWOX leads to a dose-dependent decrease in AMOTp130 levels in HEK293T cells.

### The PPxY/WW-domain interplay is important for the AMOTp130/WWOX interaction and reduced expression of AMOTp130

Here, we sought to more precisely identify the regions of AMOTp130 and WWOX that are critical for mediating this interaction. Briefly, HEK293T cells were transfected with WT WWOX and WT AMOTp130 (Fig. 2A), single PPxY (PY) motif mutants PY1, PY2 or PY3 (Fig. 2B), or with triple PY motif mutant PY123 (Fig. 2C), and an IP/Western blot assay was utilized to detect an interaction. Not surprisingly, the AMOTp130-PY123 triple mutant was unable to bind to WWOX and was undetectable on the gel (Fig. 2C, lane 2). In contrast, the single PY mutants of AMOTp130 showed varying degrees of binding to WWOX (Fig. 2B). We observed that mutant PY1 was essentially unable to interact with WWOX (Fig. 2B, lane 4), whereas both mutants PY2 and PY3 showed some degree of binding to WWOX, albeit significantly less than WT AMOTp130 (Fig. 2B, lanes 5 and 6, compare with Fig. 2A, lane 2). These data suggest that all three PY motifs of AMOTp130 participate in binding to WWOX, with PY motif #1 being most important for an efficient interaction.

**FIG. 2.**
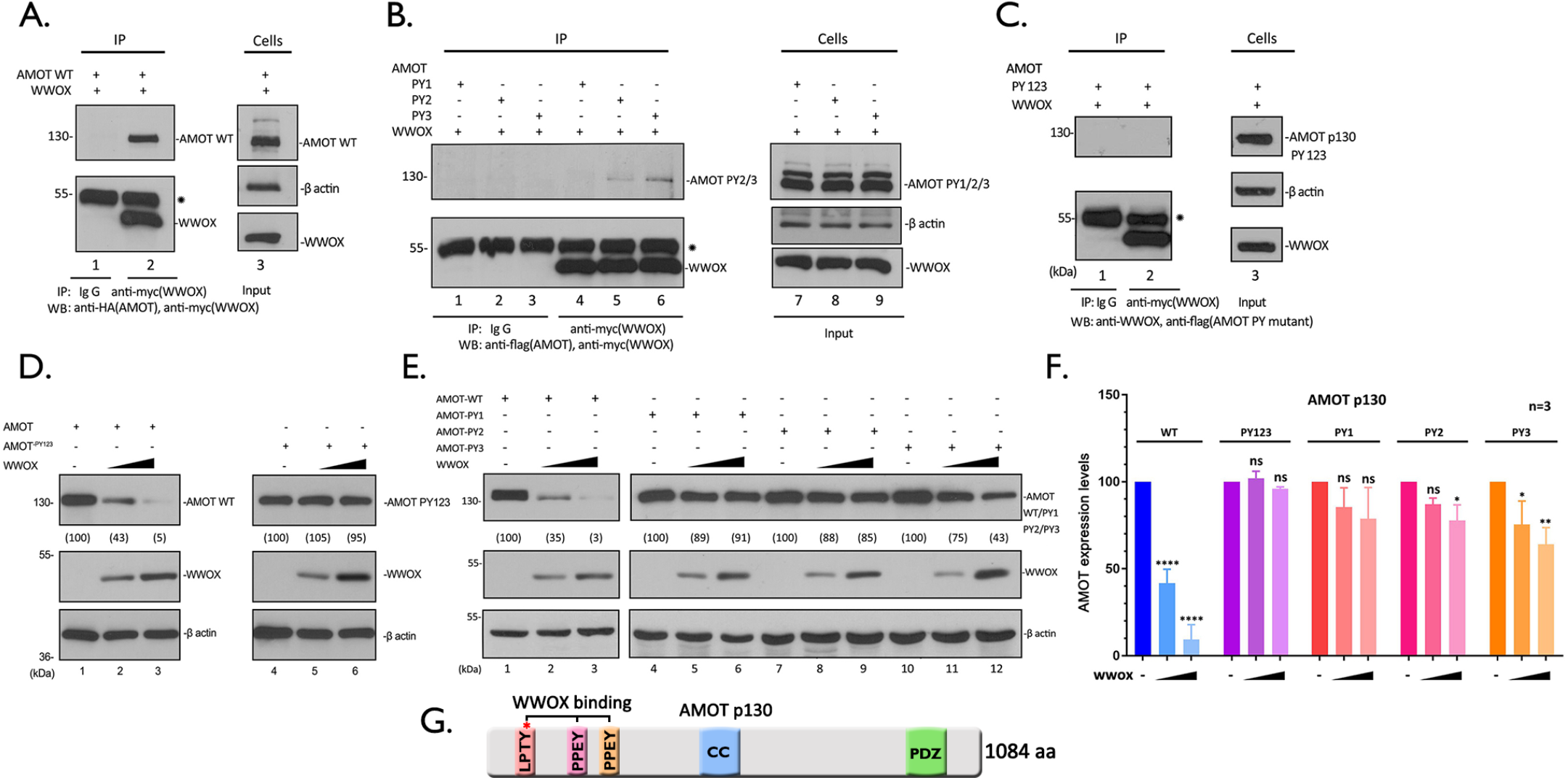
WWOX interacts with and suppresses AMOT expression in a PY motif dependent manner. **A-C)** Extracts from HEK293T cells transfected with myc-tagged WWOX and HA-tagged AMOT WT(A), flag-tagged AMOT PY1, PY2, PY3 (B) or PY123 (C) were immunoprecipitated with either normal mouse IgG or anti-myc antibody. WWOX, AMOT WT and PY motif mutants were detected in the precipitates by WB. Expression controls for AMOT WT and PY mutants, WWOX and β-actin are shown in the left panels. **D-E)** HEK293T cells were transfected with AMOT WT or PY123 (D), PY1, PY2, PY3 (E) mutants plus with vector (-) or increasing amount of WWOX, the indicated proteins were detected by WB and AMOT levels were quantified () using NIH Image-J. **F)** Quantification of the AMOT levels in (D) and (E) from three independent experiments. Statistical significance was analyzed by a one-way ANOVA. ns: not significant, *=p<0.05, **= p<0.01, ****= p<0.0001. **G)** Schematic diagram of AMOTp130 with key domains highlighted. CC = coiled coil domain; PDZ = PSD-95/Dlg1/ZO-1 domain.

Next, we wanted to ask whether increased expression of WWOX would lead to reduced levels of expression of the PY mutants of AMOTp130, as we observed for WT AMOTp130 (Figs. 1C + 1D). Briefly, HEK293T cells were co-transfected with the indicated combinations of plasmids, and protein levels were quantified by Western blotting (Figs. 2D-F). While increased expression of WWOX once again resulted in reduced levels of WT AMOTp130 (Fig. 2D, lanes 1-3), increased expression of WWOX had no effect on expression of the AMOTp130-PY123 triple mutant in repeated experiments (Fig. 2D, lanes 4-6; Fig. 2F). This finding strongly suggests that the PPxY-mediated interaction between AMOTp130 and WWOX is essential for the observed reduction in AMOTp130 expression. For the individual PY mutants, we found that increased expression of WWOX also had no significant effect on expression levels of PY1 (Fig. 2E, lanes 4-6; Fig. 2F), which correlates well with our finding that PY motif #1 is likely most critical for mediating a strong interaction with WWOX (Fig. 2B). Increased expression of WWOX had a modest effect on reducing the levels of expression of mutants PY2 and PY3 in repeated experiments (Fig. 2E, lanes 7-12; Fig. 2F).

Next, we wanted to interrogate the WW-domains of WWOX for their role in interacting with AMOTp130 and in reducing expression levels of AMOTp130 in HEK293T cells. We focused our analysis of WW-domain #1 (WW1) of WWOX, since this domain was first identified as the interacting domain with the VP40 PPxY motif and since WW-domain #2 (WW2) of WWOX is atypical in its amino acid sequence(32). Briefly, we generated a series of WW1 mutations including single point mutant Y33R, double point mutant W44A/P47A, triple point mutant Y33R/W44A/P47A, and deletion mutant ΔWW. We observed that each of the WW1 domain mutants of WWOX were reduced significantly in their ability to interact with AMOTp130 as determined by IP/Western analysis (Fig. 3A). In addition, we observed an approximate 2.5-fold and 5-fold reduction in the expression level of AMOTp130 in the presence of double mutant W44A/P47A and single mutant Y33R, respectively compared to control (Fig. 3B, lanes 3 and 4; Fig. 3C). Co-expression of mutants Y33R/W44A/P47A and ΔWW resulted in a <2-fold reduction in AMOTp130 expression compared to control (Fig. 3B, lanes 5 and 6; Fig. 3C). Thus, the more subtle mutants (Y33R and W44A/P47A) resulted in a more significant decrease in expression of AMOTp130 than that observed with the more severe mutants (Y33R/W44A/P47A, and deletion mutant ΔWW) of WWOX. Taken together, these results suggest that the PPxY/WW-domain interplay between AMOTp130 and WWOX is critical for their ability to physically interact, and plays a contributing role in the mechanism by which WWOX reduces the levels of AMOTp130 expression in HEK293T cells.

**FIG. 3.**
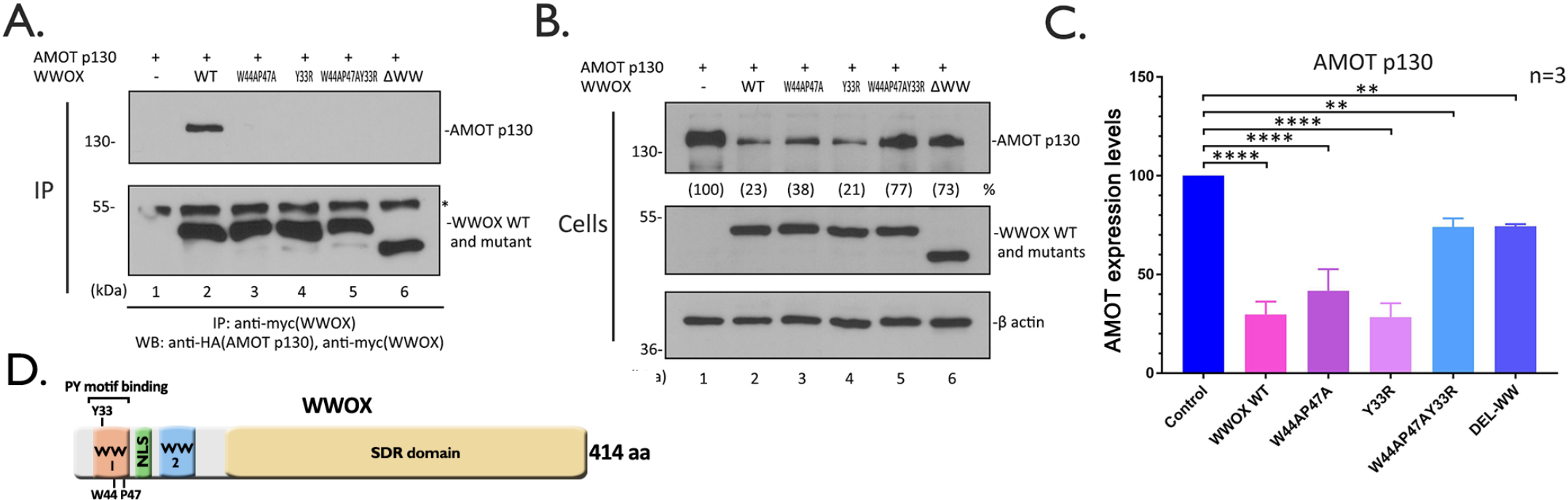
WW-domain #1 of WWOX interacts with AMOT. Extracts from HEK293T cells transfected with myc-tagged WT or the indicated mutants of WWOX and HA-tagged AMOT were subjected to IP. WWOX and AMOT in the precipitates **(A)** and cell extracts **(B)** were subjected to Western blotting analysis. **C)** Quantification of the AMOT protein levels in (B) from three independent experiments. Statistical significance was analyzed by a one-way ANOVA. **= p<0.01, ****= p<0.0001. **D)** Schematic diagram of WWOX with key domains highlighted. The Y33, W44, and P47 are three crucial amino acids in WW1 domain that mediate binding to PY motif; NLS=nuclear localization signal, SDR=Short-chain Dehydrogenase/ Reductase domain.

### WWOX alters the intracellular localization pattern of AMOTp130

We have shown previously that AMOTp130 displays a tubular, filamentous pattern of distribution in the cytoplasm of HEK293T cells as determined by confocal microscopy (31). Here, we sought to determine whether co-expression of WWOX would alter this intracellular distribution of AMOTp130, in addition to its observed role in reducing the level of AMOTp130 expression. Briefly, HEK293T cells were transfected with AMOTp130, WWOX, or both AMOTp130 + WWOX, and cells were imaged using confocal microscopy (Fig. 4). We observed the expected tubular pattern of AMOTp130 (green) throughout the cytoplasm when expressed alone, and we observed an overall diffuse cytoplasmic pattern with some punctate staining for WWOX (red) when expressed alone (Fig. 4). Interestingly, cells co-expressing AMOTp130 and WWOX revealed a dramatic change in the distribution pattern for AMOTp130 from a tubular pattern to a more punctate and perinuclear localized pattern (Fig. 4). Indeed, AMOTp130 appeared to become sequestered in small puncta/vesicles with a minimal amount of colocalization (Fig. 4, inset box) in cells expressing WWOX. In sum, we observed a profound change in the pattern of distribution for AMOTp130 in the absence vs. presence of WWOX, which likely correlates with the observed reduction in expression levels of AMOTp130 described above.

**FIG. 4.**
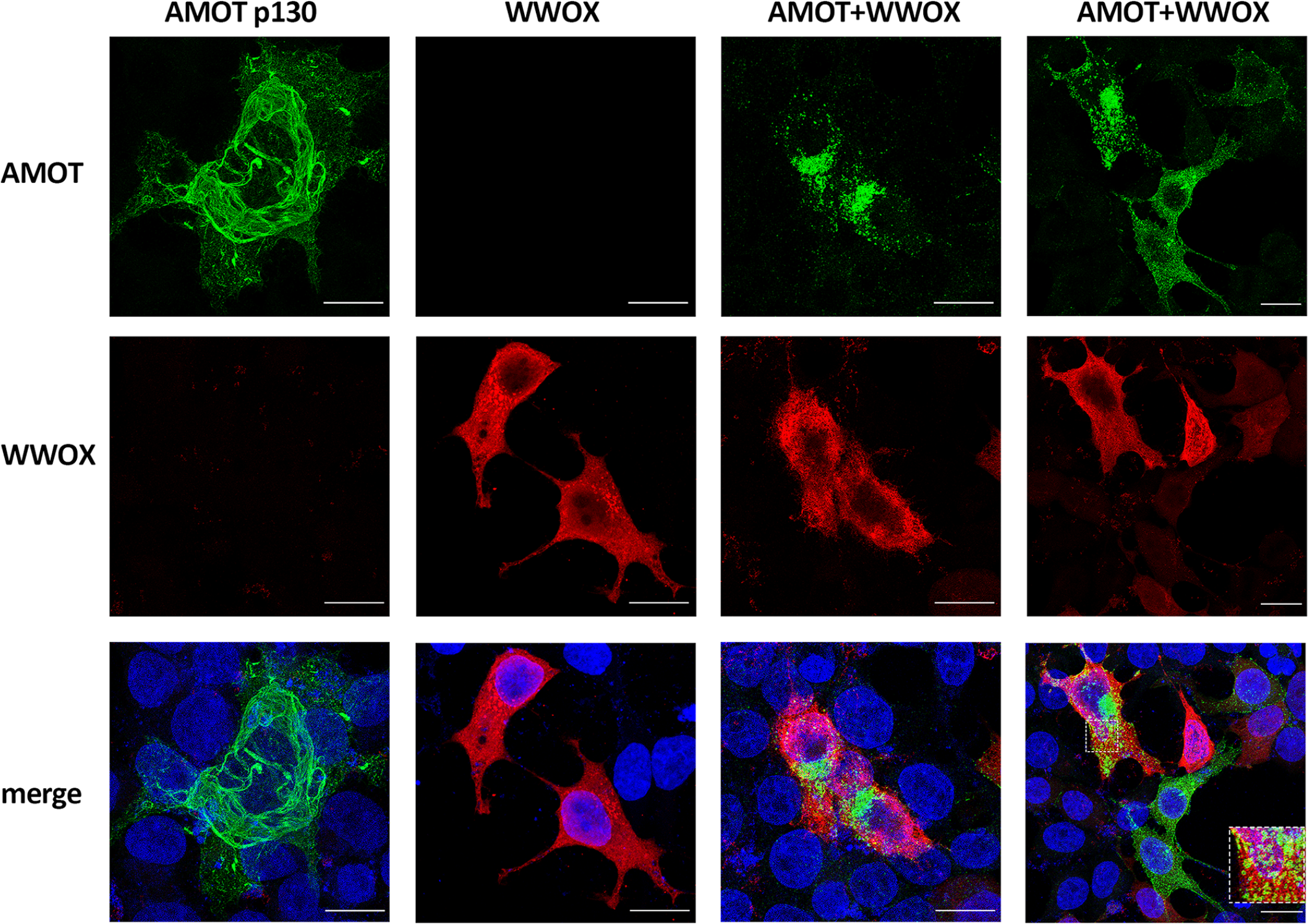
WWOX alters the intracellular distribution of AMOT. HEK293T cells were transfected with AMOT (green) and WWOX (red) alone or co-transfected with both. Cells were then visualized via immunofluorescence staining. Scale bars = 10μm.

### Increased expression of WWOX reduces AMOT expression and VP40 VLP egress

Next, we wanted to determine whether increased expression of WWOX would simultaneously reduce expression of AMOTp130 and inhibit egress of VP40 VLPs. HEK293T cells were transfected with constant amounts of AMOTp130 and eVP40 or mVP40 plasmids along with increasing amounts of WWOX, and levels of the indicated proteins were quantified in cell extracts and VLPs by Western blotting (Fig. 5). Interestingly, we observed that WWOX selectively and significantly reduced the levels of AMOTp130 in cell extracts in a dose-dependent manner without any effect on the levels of eVP40 (Figs. 5A and 5B) or mVP40 (Figs. 5C and 5D) in the same cell extracts. In contrast, the production of eVP40 (Figs. 5A and 5B) and mVP40 (Figs. 5C and 5D) VLPs was significantly decreased in a dose-dependent manner with increasing expression of WWOX (Figs. 5A and 5C, VLPs, compare lanes 2-6). These results imply that the observed WWOX-mediated reduction of AMOTp130 expression is specific, and that this reduction of AMOTp130 may negatively regulate egress of VP40 VLPs.

**FIG. 5.**
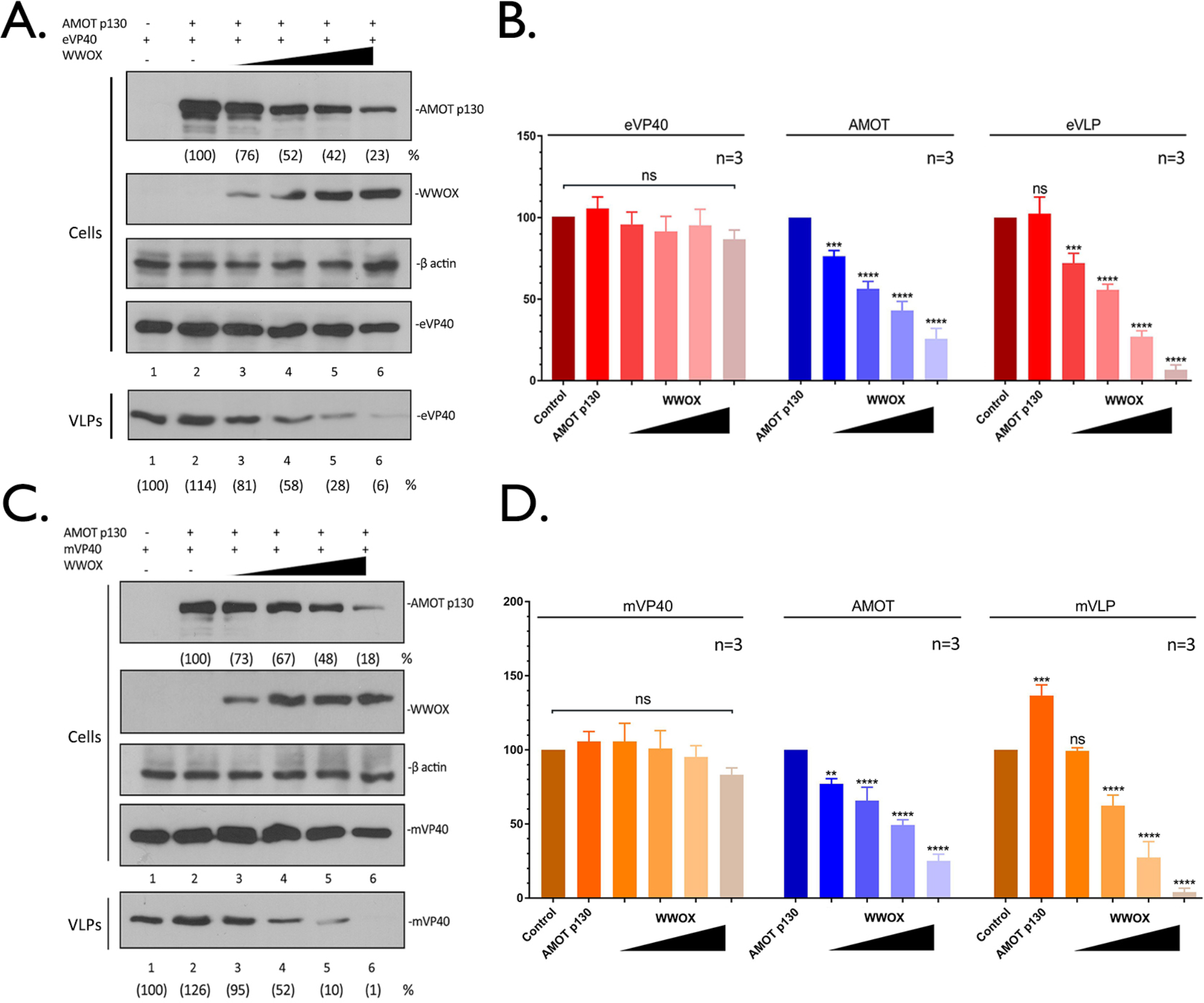
WWOX suppresses AMOT expression and inhibits VP40 VLP egress. **A and C)** HEK293T cells were transfected with a constant amount of eVP40 (A), mVP40 (C) and AMOT plus vector (-) or increasing amounts of WWOX. The indicated proteins were detected in cell extracts and VLPs by WB. The cellular levels of AMOT, eVP40, mVP40 and VP40 in VLPs were quantified using NIH Image-J. The amounts of eVP40 (A, cells, lane 1), mVP40 (C, cells, lane 1), AMOT (A, C cells, lane 2) in control cells were set at 100%. Also, eVP40 (A, VLPs, lane 1) and mVP40 (C, VLPs, lane 1) VLP production from control cells was set at 100%. Numbers in () represent relative protein levels and VLP budding efficiency compared to the control. **B and D)** Quantification of the indicated cellular protein levels and relative budding efficiency of eVP40 (B) and mVP40 (D) VLPs from three independent experiments. Statistical significance was analyzed by a one-way ANOVA. ns: not significant, **=p<0.01, ***=p<0.001, ****= p<0.0001.

### WWOX and AMOT affect the intracellular localization of VP40

Next, we used confocal microscopy to visualize any changes in the spatial distribution of VP40 in the absence or presence of WWOX and AMOTp130. Briefly, eVP40 and AMOT were expressed in HEK 293T cells in the absence (Fig. 6, top row) or presence (Fig. 6, middle and bottom row) of WWOX, and representative confocal images are shown. As expected, eVP40 was present throughout the cytoplasm with light accumulation at the plasma membrane, and AMOT was distributed as tubular bundles throughout the cells (Fig. 6, top row, arrow). Conversely, VP40 appeared to accumulate more heavily at the cell periphery and could be detected more readily in the nucleus in cells co-expressing both AMOT and WWOX (Fig. 6, middle and bottom rows, triangles). Notably, AMOT no longer displayed the tubular pattern in these cells, but rather was more punctate and disperse. These data show that exogenous expression of WWOX resulted in redistribution of both AMOT and VP40. It is tempting to speculate that WWOX disrupts the normal cytoskeletal association of AMOT leading to its degradation in punctate vesicles. Moreover, WWOX may “drag” a portion of VP40 into the nucleus and the remainder of VP40 accumulates at the plasma membrane where it remains tethered and unable to bud efficiently due in part to the disruption of AMOT tubular distribution (see Supp. Video S1).

**FIG. 6.**
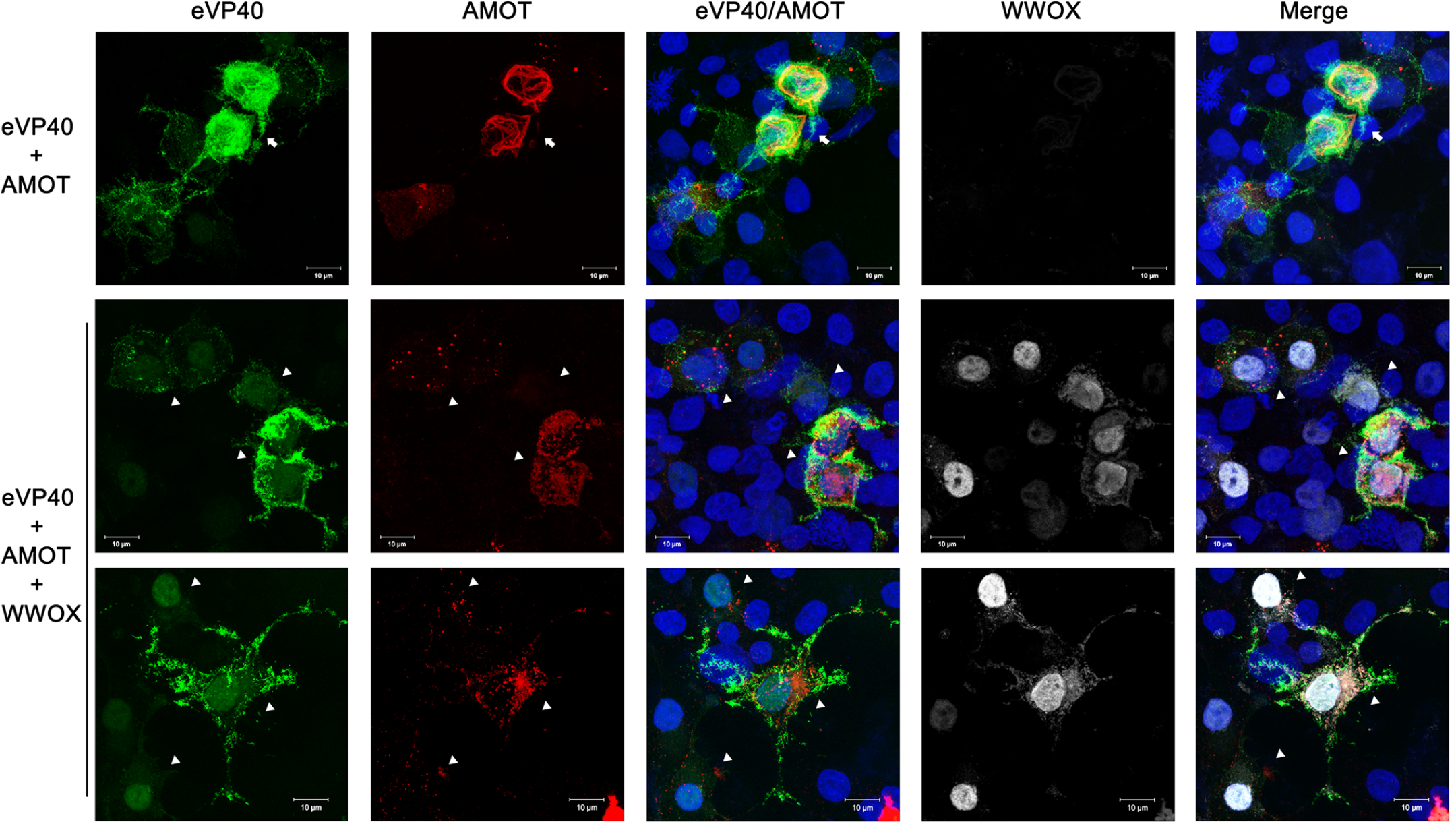
Expression of WWOX and AMOT affects localization of VP40. HEK293T cells were transfected with eVP40 (green) and AMOT (red) alone, or with WWOX (white) and visualized by confocal microscopy. Scale bars = 10µm.

### Inhibition of VP40 VLP egress by WWOX is dependent on expression of AMOTp130

To further test our hypothesis that disruption of AMOTp130 expression by WWOX leads to a decrease in VP40 VLP egress, we used shCtrl and shAMOT knockdown cell lines in our VLP budding assay (Fig. 7). Briefly, shCtrl and shAMOT cells were transfected with a constant amount of eVP40 (Figs. 7A + 7B) or mVP40 (Figs. 7C + 7D) in the absence or presence of increasing amounts of WWOX, and cell lysates and VLPs were harvested for analysis by Western blotting. We observed a significant decrease in both eVP40 (10-fold) and mVP40 (50-fold) VLP egress in the shCtrl cells in the presence of WWOX (Figs. 7A and 7C, compare lanes 1-3). As expected from prior results, budding of both eVP40 and mVP40 VLPs is significantly reduced in shAMOT cells compared to that in shCtrl cells (Figs. 7A and 7C, compare lanes 1 and 4). Notably, expression of WWOX did not significantly decrease budding of either eVP40 (<2-fold) or mVP40 (2-fold) in the shAMOT cells (Figs. 7A and 7C, compare lanes 4-6). In sum, these results support our hypothesis that inhibition of VP40 VLP egress by WWOX is due, in part, to WWOX’s physical interaction with, and functional disruption of, AMOTp130.

**FIG. 7.**
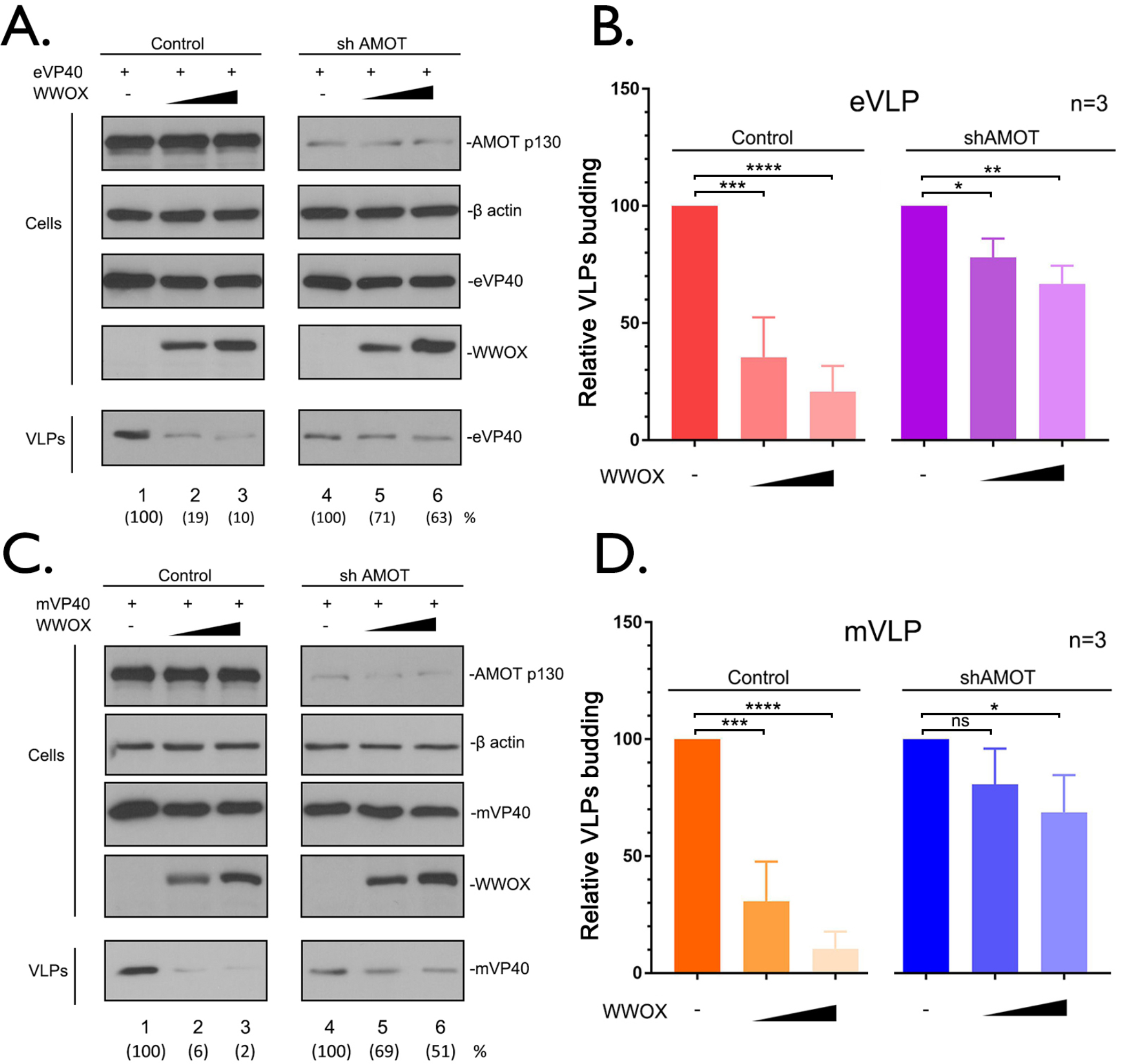
Expression of AMOT is required for WWOX-mediated inhibition of VP40 VLP egress. **A and C)** shCtrl and shAMOT cells were transfected with a constant amount of eVP40 (A) or mVP40 (B) with vector alone (-) or increasing amounts of WWOX. The indicated proteins were detected in cell extracts and VLPs by WB. VP40 levels in VLPs were quantified () using NIH Image-J software. **B and D)** Quantification of the relative budding efficiency of eVP40 (B) or mVP40 (D) VLPs from three independent experiments (n=3). WWOX minus samples were normalized independently for Control and shAMot conditions (B and D). Statistical significance was analyzed by a one-way ANOVA. ns: not significant, *=p<0.05, **=p<0.001, ***=p<0.001, ****= p<0.0001.

### WWOX induces lysosomal degradation of AMOTp130

The ubiquitin-proteasome system and the lysosomal pathway are the two major pathways involved in protein degradation and turnover in eukaryotic cells. As a multifunctional scaffolding protein, AMOTp130 expression and stability are tightly regulated by other host proteins/pathways including LATS1/2 kinases and E3 ubiquitin ligases such as Nedd4 and Itch (25, 33–39). Thus, it was of interest to determine more precisely how WWOX expression may induce degradation of AMOTp130. Since AMOTp130 appeared to be sequestrated in punctate vesicles in the presence of WWOX (Figs. 4 and 6), we hypothesize that AMOTp130 may be subjected to degradation by the lysosomal pathway. To test our hypothesis, we utilized confocal microscopy and incorporated lysosomal marker protein LAMP1, to determine whether AMOTp130 localizes to lysosomes in the presence of WWOX (Fig. 8A). We observed the expected tubular pattern of AMOTp130 in the absence of WWOX, and under these conditions, AMOTp130 did not colocalize with LAMP1 (Fig. 8A, top row). In contrast, we observed clear colocalization of AMOTp130 in vesicles containing LAMP1 and the loss of the tubular pattern in the presence of WWOX (Fig. 8A, bottom row, arrows and zoomed view).

**FIG. 8.**
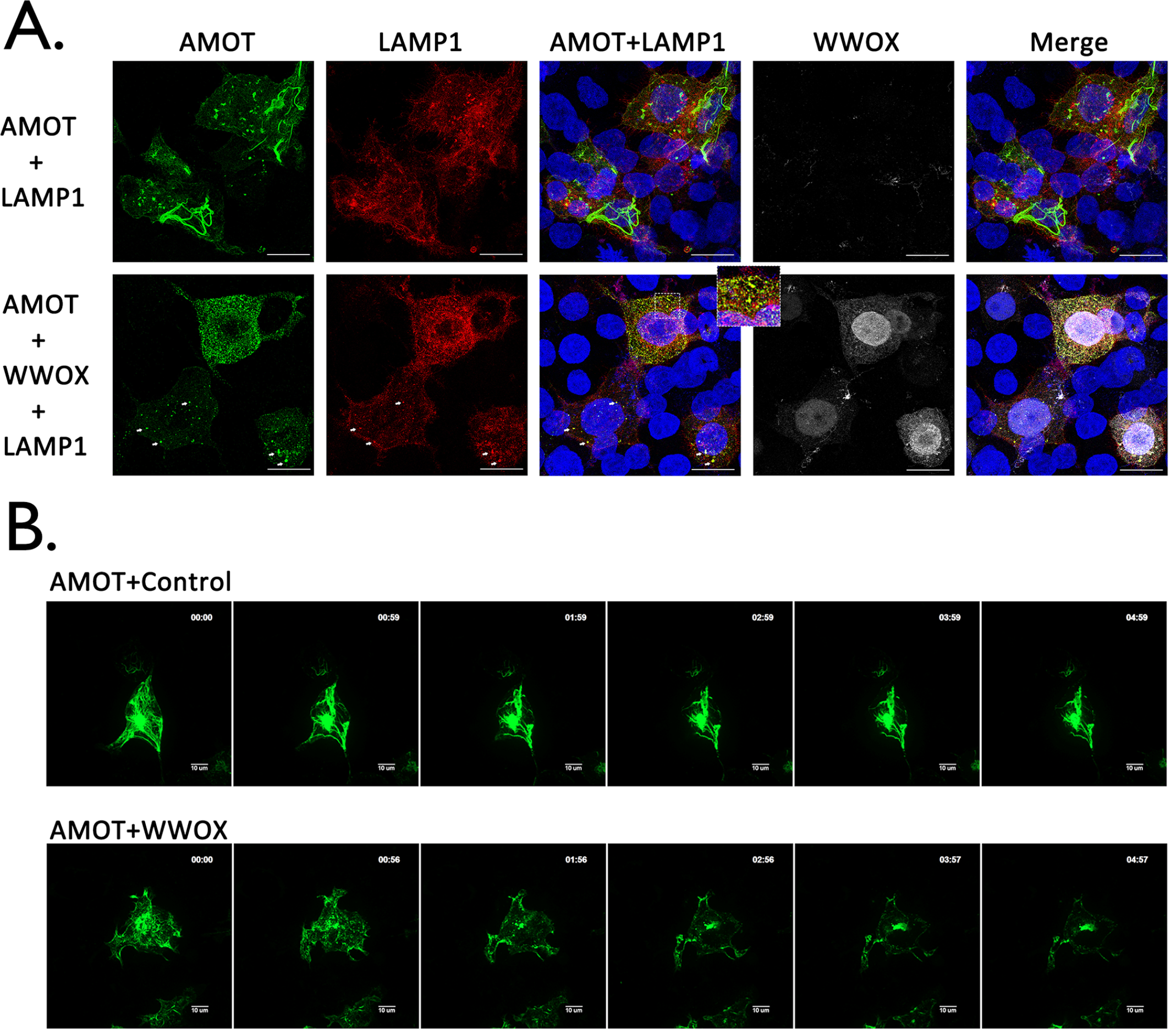
WWOX induces lysosomal degradation of AMOT. **A)** HEK293T cells were transfected with the indicated combinations of plasmids including lysosome marker mCherry-LAMP1 (red), AMOT (green), and WWOX (white). Cells were visualized using confocal microscopy. Scale bars = 10μm. The zoomed view and arrows highlight co-localization of AMOT and LAMP1 in lysosomes. **B)** HEK293T cells were transfected with YFP-AMOT and mCherry-LAMP1 plus vector (Control) or WWOX. Live cells were observed via spinning disk confocal microscopy beginning at 12 hours post transfection. Representative images showing the localization of AMOT in control and WWOX expressing cells at each hour during observation. Scale bars = 10μm.

To further illustrate the temporal and spatial distribution dynamics of AMOTp130 and LAMP1 in the absence or presence of WWOX, we conducted time-lapse confocal microscopy using live HEK293T cells (Fig. 8B). Briefly, YFP-AMOTp130 and mCherry-LAMP1 fusion proteins were co-expressed in HEK 293T cells with vector alone or WWOX. At 12 hours post-transfection, cells were subjected to live cell spinning disk confocal microscopy for 5 hours, and images were taken every 10 minutes. Representative images at each hour time point are shown highlighting the changes in localization of AMOTp130 in the absence or presence of WWOX (Fig. 8B). We observed that the tubular pattern of AMOTp130 did not change significantly over time in cells lacking WWOX (Fig. 8B, top panels). In contrast, the tubular distribution pattern of AMOTp130 was altered to a more punctate pattern over time in the presence of WWOX (Fig. 8B, bottom panels). Taken together, these results suggest that the mechanism by which WWOX reduces expression of AMOTp130 involves lysosomal-mediated degradation.

### Pharmacological inhibition of lysosome function restores expression of AMOTp130 and rescues VP40 VLP budding in the presence of WWOX

If WWOX-mediated degradation of AMOTp130 occurs via the lysosomal pathway, then we reasoned that treating cells with lysosomal inhibitor chloroquine (CQ) (40, 41) should restore expression of AMOTp130 and rescue VP40 VLP egress in the presence of WWOX. To assess restoration of AMOTp130 expression, HEK293T cells were either mock-treated, or treated with CQ and transfected with the indicated combination of AMOTp130 and WWOX plasmids (Figs. 9A and 9B). As expected, AMOTp130 levels were significantly reduced in mock-treated cells in the presence of WWOX (Fig. 9A, lanes 1-3; Fig. 9B); however, there was no significant change in AMOTp130 levels in cells treated with CQ in the presence of WWOX (Fig. 9A, lanes 4-6; Fig. 9B). When we co-expressed eVP40 under the same conditions, we observed almost a complete rescue of eVP40 VLP budding back to WT levels in cells treated with CQ in the presence of WWOX (Fig. 9C, lanes 5 and 6; Fig. 9D) compared to mock-treated controls (Fig. 9C, lanes 3 and 4; Fig. 9D). These findings suggest that the rescue of eVP40 VLP egress in CQ-treated cells is likely due, in part, to the restoration of AMOTp130 back to WT levels.

**FIG. 9.**
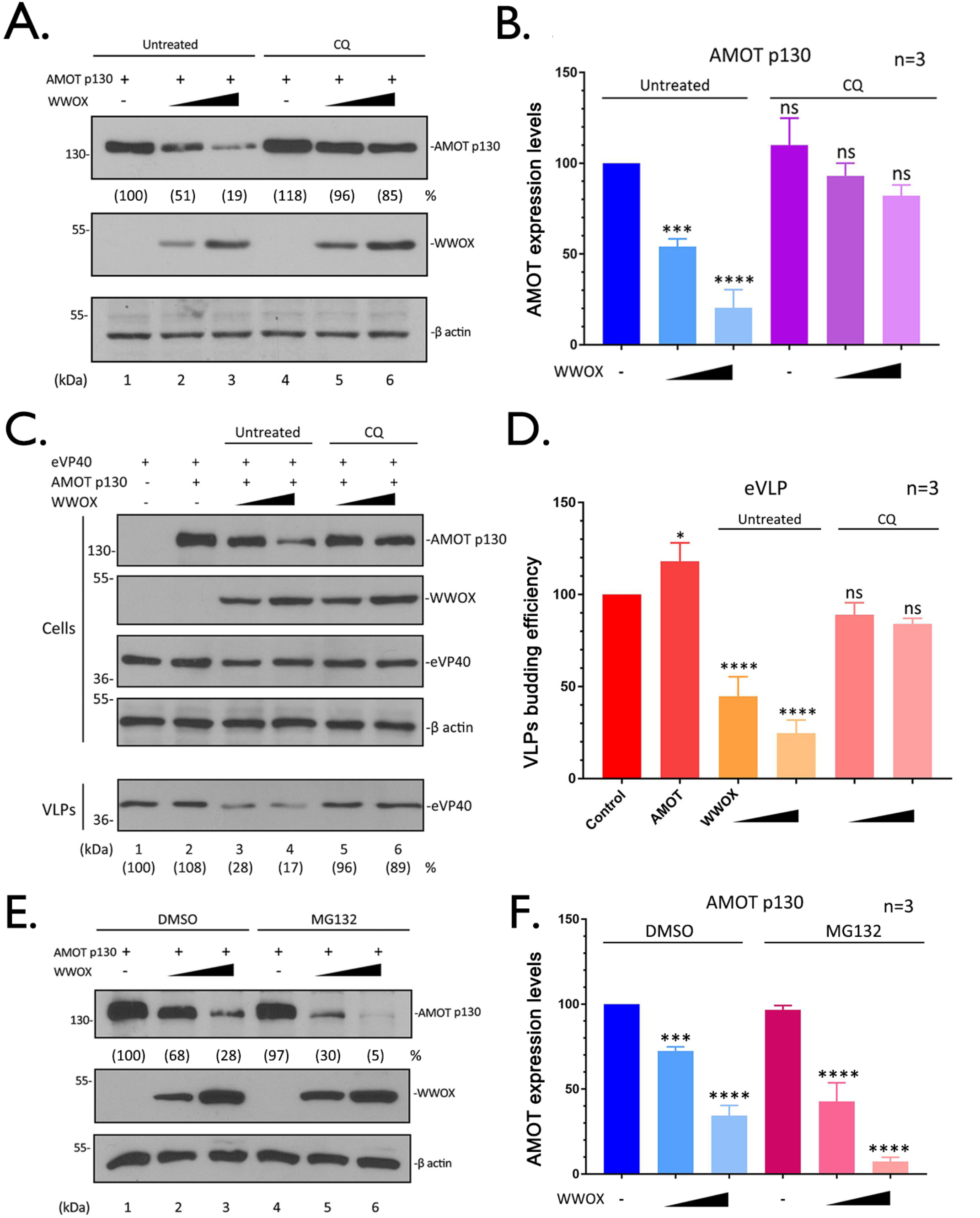
Pharmacological inhibition of lysosome function restores AMOT expression and rescues VP40 VLP budding. **A)** HEK293T cells were transfected with a constant amount of AMOT plus vector (-) or increasing amounts of WWOX. After transfection, cells were treated with (lanes 4-6) or without (lanes 1-3) lysosomal inhibitor chloroquine (50µM) for 16 hours. The indicated proteins were detected in cell extracts by WB. AMOT levels were quantified () using NIH Image-J. **B)** Quantification of AMOT (A) from three independent experiments (n=3). Statistical significance was analyzed by a one-way ANOVA. ns: not significant, ***=p<0.001, ****= p<0.0001. **C)** HEK293T cells were transfected with a constant amount of eVP40, and AMOT plus vector (-) or increasing amounts of WWOX. Cells were treated with (lanes 5, 6) or without (lanes 3, 4) CQ (50µM) for 16 hours. The indicated proteins were detected in cell extracts and VLPs by WB. The yields of eVP40 VLPs were quantified () using NIH Image-J software. **D)** Quantification of the relative budding efficiency of eVP40 VLPs under the indicated conditions from three independent experiments (n=3). Statistical significance was analyzed by a one-way ANOVA. ns: not significant, *=p<0.05, ****= p<0.0001. **E)** HEK293T cells were transfected with the a constant amount of AMOT plus vector (-) or increasing amounts of WWOX. Cells were treated with (lanes 4-6) or without (lanes 1-3) proteasomal inhibitor MG132 (10µM) for 8 hours before harvesting. The indicated proteins were detected in cell extracts by WB. **F)** Quantification of AMOT (E) from three independent experiments (n=3). Statistical significance was analyzed by a one-way ANOVA. ns: not significant, ***=p<0.001, ****= p<0.0001.

Lastly, we asked whether the ubiquitin-proteasome system may also be involved in WWOX-mediated degradation of AMOTp130 using a pharmacological approach. Briefly, HEK293T cells were treated with DMSO or the proteasome inhibitor, MG132 (39), and cells were transfected with the indicated combinations of plasmids (Fig. 9E). In contrast to our findings following treatment with CQ, treatment of cells with MG132 did not lead to restoration of AMOTp130 levels in the presence of WWOX (Fig. 9E, 9F). In sum, these results suggest that WWOX represses AMOT by induction of its lysosomal degradation, and thus modulates AMOT and eventually leads to the inhibition of VLPs egress.

## DISCUSSION

EBOV and MARV VP40 matrix protein utilizes L-domain motifs (PPxY, PTAP, YxxL) to recruit specific host proteins to facilitate virus egress and dissemination (5, 8, 9, 42, 43). In addition to the recruitment of host proteins that positively regulate budding (*e.g.* Tsg101, Nedd4, Itch, WWP1, and Smurf2) (11, 16, 17, 42, 44), the VP40 PPxY L-domain also engages with host proteins that negatively regulate VP40-mediated egress (*e.g.* BAG3, YAP/TAZ, and WWOX) (12, 13, 15). Thus, the modular and competitive nature of the PPxY-WW domain interplay likely impacts both host and virus functions (18). Notably, host PPxY/WW-domain interactions regulate diverse signaling networks and major cellular processes, such as the Hippo pathway and cell division/migration.

Here, we describe how the physical and functional interaction between host AMOTp130 and host WWOX affects VP40 VLP budding (Fig. 10). AMOTp130 is a key multi-PPxY containing host protein that engages host WW-domain containing proteins that both positively and negatively impact viral budding. For example, AMOTp130 interacts with the YAP, BAG3, and WWOX in a PPxY/WW-domain dependent manner to function as a “master regulator” of several physiologically relevant pathways/processes, including transcription (Hippo pathway), apoptosis, cytoskeletal dynamics, and tight junction (TJ) integrity (19, 23, 25, 26, 28, 45–48). In addition, AMOT stability and turnover is tightly regulated via PPxY/WW-domain interactions with Nedd4 E3 ubiquitin ligase family members (34–36, 39). While Amot was previously shown to regulate assembly and egress of non-PPxY-containing viruses (49, 50), we recently revealed a role for endogenous AMOT in positively regulating egress of PPxY-containing EBOV and MARV (12, 13, 31).

**FIG. 10.**
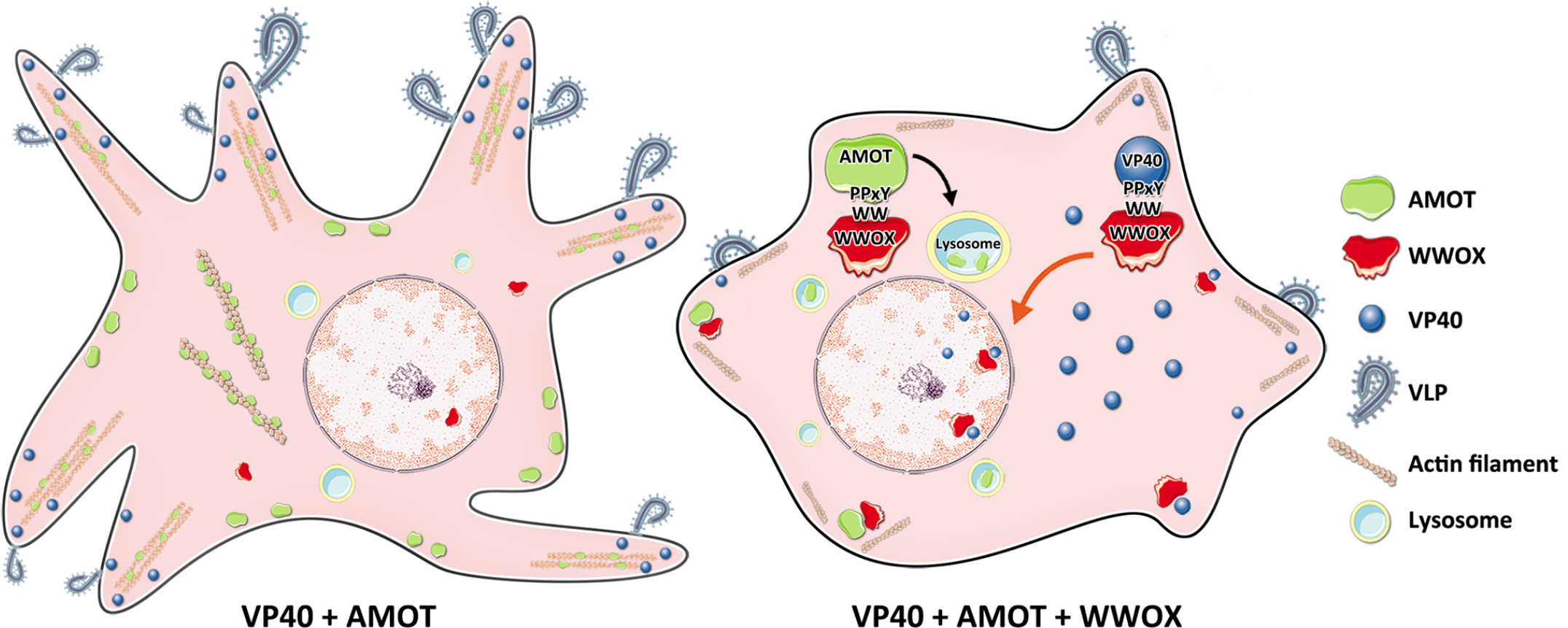
Working model of PPxY/WW-domain interactions among AMOT, WWOX and VP40. **Left:** AMOT facilitates VP40 VLP egress via its ability to bind actin and regulate its dynamics at the plasma membrane. **Right:** Exogenously expressed WWOX interacts with both AMOT and VP40 in a PPxY/WW-domain dependent manner. These interactions lead to reduced levels of VP40 at the plasma membrane, enhanced nuclear localization of VP40 (red arrow), and lysosomal mediated degradation of AMOT (black arrow); all of which results in a decrease of VLP egress.

We showed that WWOX reduced expression and modulated intracellular distribution of AMOTp130 in a PPxY/WW-domain dependent manner. Indeed, we observed a dose-dependent reduction in expression of PPxY-containing AMOTp130, but not PPxY-lacking AMOTp80, in the presence of WWOX (Fig. 1). Moreover, expression of a mutant AMOTp130 containing mutations in all three PPxY motifs was not affected by exogenous expression of WWOX (Fig. 2), and a WW-domain mutant of WWOX did not reduce the levels of AMOTp130 compared to that of WT WWOX (Fig. 3).

Under conditions of exogenous expression of WWOX, we observed that budding of both eVP40 and mVP40 VLPs was significantly reduced (Fig. 4). We hypothesized that one possible mechanism could be that WWOX was negatively regulating budding of VP40 VLPs indirectly, by reducing expression of AMOTp130 and preventing it from facilitating egress of VLPs from the plasma membrane as reported previously (13, 31). In support of this hypothesis, we found that budding of both eVP40 and mVP40 VLPs was not significantly reduced in shAMOT cells in the presence of WWOX, but was significantly reduced in shCtrl cells in the presence of WWOX (Fig. 7). These findings suggest that WWOX’s negative effect on VP40 VLP egress is the result of a novel, indirect mechanism of action that requires expression of, and likely an interaction with, endogenous AMOTp130.

We sought to further understand the mechanism by which WWOX reduced expression of AMOTp130. Toward this end, our results suggest that WWOX mediates reduction of AMOTp130 levels via the lysosomal degradation pathway as judged by confocal microscopy and the use of pharmacological inhibitors (Figs. 8 and 9). Notably, WWOX was unable to reduce expression of AMOTp130 in cells treated with chloroquine compared to controls. These conditions also resulted in the restoration of budding of VP40 VLPs to near WT levels in cells expressing WWOX and treated with chloroquine. In contrast to the results with chloroquine, WWOX retained the ability to reduce the levels of AMOTp130 in cells treated with MG132, suggesting that WWOX-mediated degradation of AMOTp130 was not occurring via the ubiquitin/proteasome pathway.

In addition to the indirect mechanism of inhibition of VLP budding described above, our data also suggest that WWOX can inhibit egress of VP40 VLPs via a direct PPxY/WW-domain interaction that leads to reduced levels of VP40 at the plasma membrane and increased levels of VP40 detected in the nucleus (Fig. 10) (12). Indeed, we observed enhanced nuclear localization of VP40 in cells expressing WWOX. Since WWOX contains a nuclear localization signal and normally shuttles in and out of the nucleus (51), one possibility is that WWOX may sequester or drag some VP40 into the nucleus as a result of a direct PPxY/WW-domain interaction, leading to a subsequent decrease in VLP budding. Interestingly, nuclear localization of eVP40 has been reported previously (52–54); however, a functional role for nuclear eVP40 has not been described. Investigations into a potential role for eVP40 in the nucleus may be warranted due in part to the identification of nuclear transcriptional regulators such as WWOX and YAP/TAZ as specific host interactors. It is tempting to speculate that host proteins such as WWOX and YAP/TAZ may interact with and translocate eVP40 into the nucleus where these virus-host complexes may then affect transcription of WWOX and/or YAP/TAZ responsive genes to generate a cellular environment beneficial for virus replication, budding, and/or disease progression. Alternatively, since WWOX is known to directly bind to multiple transcriptional activators, such as p38, p73, AP-2γ, ErBb4, c-Jun and RUNX2, in a PPxY/WW domain dependent manner (22, 23, 55–59), it will be of interest to determine whether there is any competitive interplay among these host proteins and PPxY-containing VP40 proteins in VP40 expressing or virus infected cells that may result in a biological consequences having an impact on both the virus and host. Such studies may also provide novel insights into the development of new host-oriented antivirals that target these modular virus-host interactions.

## MATERIALS AND METHODS

### Cells, antibodies, and plasmids

HEK293T-based shCtrl and shAMOT cells (kindly provided by J. Kissil, Scripps Research, FL) and HEK293T cells were maintained in Dulbecco’s modified Eagle’s medium (DMEM) (CORNING) supplemented with 10% fetal bovine serum (FBS) (GIBCO), penicillin (100U/ml)/streptomycin (100μg/ml) (INVITROGEN). Cells were grown at 37°C in a humidified 5% CO_2_ incubator. The primary antibodies used in this study include mouse anti-myc (Millipore), mouse anti-HA antibody (Sigma), mouse anti-flag (Sigma), mouse anti-AMOT (Santa Cruz), rabbit anti-eVP40 (IBT), mouse anti β-actin (Proteintech). The plasmids encoding eVP40, GFP-eVP40 were described previously (8, 42). Flag-tagged mVP40 was kindly provided by S. Becker (Institut für Virologie, Marburg, Germany). The flag tagged AMOT PY123, PY1, PY2 and PY3 mutants were kindly provided by J. Kissil (Scripps Research Institute, FL). The YFP-tagged AMOTp130 was kindly provided by K-L. Guan (University of California, San Diego). The mCherry-tagged LAMP1 was a gift from Amy Palmer (Addgene plasmid # 45147). The myc-tagged WWOX plasmid was kindly provided by R. I. Aqeilan (Jerusalem, Israel). The mutants of WWOX were generated via QuikChange™ method, and primers used are as follows:

<u>Y33R1</u>: 5’GGTGTGATTGGCGTAGCGAACCCAGCCGTCCTTG3’,

<u>Y33R2</u>: 5’CAAGGACGGCTGGGTTCGCTACGCCAATCACACC3’.

<u>W44AP47A1</u>:5’TTTCTTTTTCCAGTTTTTGCATGTTCCGCCTGAGTCTTCTCCTCGGTG3’,

<u>W44AP47A2</u>:5’CACCGAGGAGAAGACTCAGGCGGAACATGCAAAAACTGGAAAAAGAAA3’.

<u>ΔWW1</u>: 5’CATCCACAGTAAACGCGTCCTCACTGTCCGTG3’,

<u>ΔWW2</u>: 5’CACGGACAGTGAGGACGCGTTTACTGTGGATG3’.

### Immunoprecipitation assay

HEK293T cells seeded in 6 well plates were transfected with the indicated plasmid combinations using Lipofectamine reagent (INVITROGEN). At 24 hours post transfection, cells were harvested and lysed, and the cell extracts were subjected to Western blotting (WB) and co-immunoprecipitation (IP). The protein complexes were precipitated by either mouse IgG or anti-myc antibody. First, the cell extracts were incubated with antisera overnight at 4°C with continuous rotation, then the protein A/G agarose beads (Santa Cruz) were added to the mixtures and incubated for 5 hours with continuous rotation. After incubation, beads were collected via centrifugation and washed 5 times. The input cell extracts and immunoprecipitates were then detected by WB with appropriate antisera as indicated.

### Western blotting and VLP budding assays

HEK293T cells were transfected with 0.25μg AMOT p130 or p80 plus with increasing amounts (0.1, 0.25, 0.5, 1.0μg) of WWOX plasmids, or cells were transfected with 0.25μg AMOTp130 WT or PY123, PY1, PY2, PY3 mutants plus with increasing amounts (0.25, 0.5μg) of WWOX plasmids. The total amount of transfected DNA was equivalent in all samples. Cell extracts were harvested at 24 hours post transfection then subjected to SDS-PAGE and WB analyses.

HEK293T cells were transfected with 0.25μg AMOT and 0.5μg WT or mutant WWOX. Cell extracts were subjected to WB and IP analyses. For VLP budding and WWOX titration experiments, HEK293T cells were transfected with 0.2μg of eVP40 or flag-tagged mVP40, plus 0.25μg AMOTp130 and increasing amounts (0.1, 0.25, 0.5, 1.0μg) of WWOX plasmids. The eVP40 and mVP40 in VLPs and the indicated proteins in cell extracts were detected by WB.

### Indirect immunofluorescence assay

HEK293T cells were transfected with the indicated plasmid combinations. At 24 hours post transfection, cells were washed with cold PBS and fixed with 4% formaldehyde for 20 min at room temperature, then permeabilized with 0.2% Triton X-100. After washing 3X with PBS, cells were blocked for 1 hour, then incubated with rabbit anti-myc (WWOX) or mouse anti-HA (AMOT) antisera. Next, cells were stained with Alexa Fluor 488, 594 or 647 goat anti-mouse/rabbit secondary antibodies (LIFE TECHONOLOGIES). The GFP-eVP40 and mCherry-LAMP1 were visualized via fluorescent tag. Cells were mounted with ProLong™ Glass Antifade Mountant with Hoechst 33342 (LIFE TECHONOLOGIES). Microscopy was performed using a Leica SP5 FLIM inverted confocal microscope. Serial optical planes of focus were taken, and the collected images were merged into one by using the Leica microsystems (LAS AF) software.

### VLP budding assay in HEK293T shCtrl and shAMOT cells

HEK293T shCtrl and shAMOT cells were transfected with 0.2μg of eVP40 or mVP40 plus vector or 0.25, 0.5μg of WWOX. VP40 VLPs and eVP40, mVP40, WWOX and endogenous AMOTp130 in cell extracts were detected by WB using appropriate antisera.

### Live cell imaging and time-lapse microscopy

HEK293T cells were seed on chambered coverglasses and transfected with YFP-AMOT (0.25μg) and mCherry-LAMP1 (0.25μg) plus vector or WWOX (0.5μg). Live cells were observed at 12 hours post transfection using a Leica DMI4000 microscope with Yokagawa CSU-X1 spinning disk confocal attachment. Images were taken every 10 minutes over a 5-hour window of observation.

### Pharmacological inhibition of lysosome or proteosome functions

HEK293T cells were transfected with 0.25μg AMOTp130 plus vector or 0.25, 0.5μg WWOX plasmids. For lysosomal inhibition, cells were untreated or treated with CQ (50μM) for 16 hours. For proteasomal inhibition, cells were treated with DMSO or MG132 (10μM) at 8 hours before harvest. The indicated proteins were detected via WB. For VLP budding, HEK293T cells were transfected with 0.2μg eVP40 alone or with 0.25μg AMOTp130 and 0.25, 0.5μg WWOX plasmids. At 6 hours post transfection, cells were untreated or treated with CQ (50μM) for 16 hours. Then cell extracts and VLPs were harvested and subjected to WB analysis.

## ACKNOWLEDGEMENTS

The authors would like to thank J. Kissil, R. Aqeilan, K-L Guan, and M. Sudol for kindly providing reagents. Funding was provided in part by National Institutes of Health grants AI138052, AI139392, AI153815, and EY031465 to RNH.

## REFERENCES

1. Malvy D, McElroy AK, de Clerck H, Günther S, van Griensven J. 2019. Ebola virus disease. The Lancet 393:936–948.

2. Sweileh WM. 2017. Global research trends of World Health Organization’s top eight emerging pathogens. Global Health 13:9.

3. Nyakarahuka L, Shoemaker TR, Balinandi S, Chemos G, Kwesiga B, Mulei S, Kyondo J, Tumusiime A, Kofman A, Masiira B, Whitmer S, Brown S, Cannon D, Chiang CF, Graziano J, Morales-Betoulle M, Patel K, Zufan S, Komakech I, Natseri N, Chepkwurui PM, Lubwama B, Okiria J, Kayiwa J, Nkonwa IH, Eyu P, Nakiire L, Okarikod EC, Cheptoyek L, Wangila BE, Wanje M, Tusiime P, Bulage L, Mwebesa HG, Ario AR, Makumbi I, Nakinsige A, Muruta A, Nanyunja M, Homsy J, Zhu BP, Nelson L, Kaleebu P, Rollin PE, Nichol ST, Klena JD, Lutwama JJ. 2019. Marburg virus disease outbreak in Kween District Uganda, 2017: Epidemiological and laboratory findings. PLoS Negl Trop Dis 13:e0007257.

4. Makino A, Yamayoshi S, Shinya K, Noda T, Kawaoka Y. 2011. Identification of amino acids in Marburg virus VP40 that are important for virus-like particle budding. J Infect Dis 204 Suppl 3:S871–7.

5. Liu Y, Cocka L, Okumura A, Zhang YA, Sunyer JO, Harty RN. 2010. Conserved motifs within Ebola and Marburg virus VP40 proteins are important for stability, localization, and subsequent budding of virus-like particles. J Virol 84:2294–303.

6. Kolesnikova L, Ryabchikova E, Shestopalov A, Becker S. 2007. Basolateral budding of Marburg virus: VP40 retargets viral glycoprotein GP to the basolateral surface. J Infect Dis 196 Suppl 2:S232–6.

7. Kolesnikova L, Bugany H, Klenk HD, Becker S. 2002. VP40, the matrix protein of Marburg virus, is associated with membranes of the late endosomal compartment. J Virol 76:1825–38.

8. Harty RN, Brown ME, Wang G, Huibregtse J, Hayes FP. 2000. A PPxY motif within the VP40 protein of Ebola virus interacts physically and functionally with a ubiquitin ligase: implications for filovirus budding. Proc Natl Acad Sci U S A 97:13871–6.

9. Harty RN. 2009. No exit: targeting the budding process to inhibit filovirus replication. Antiviral Res 81:189–97.

10. Harty RN. 2018. Hemorrhagic Fever Virus Budding Studies. Methods Mol Biol 1604:209–215.

11. Shepley-McTaggart A, Schwoerer MP, Sagum CA, Bedford MT, Jaladanki CK, Fan H, Cassel J, Harty RN. 2021. Ubiquitin Ligase SMURF2 Interacts with Filovirus VP40 and Promotes Egress of VP40 VLPs. Viruses 13.

12. Liang J, Ruthel G, Sagum CA, Bedford MT, Sidhu SS, Sudol M, Jaladanki CK, Fan H, Freedman BD, Harty RN. 2021. Angiomotin Counteracts the Negative Regulatory Effect of Host WWOX on Viral PPxY-Mediated Egress. J Virol 95.

13. Han Z, Dash S, Sagum CA, Ruthel G, Jaladanki CK, Berry CT, Schwoerer MP, Harty NM, Freedman BD, Bedford MT, Fan H, Sidhu SS, Sudol M, Shtanko O, Harty RN. 2020. Modular mimicry and engagement of the Hippo pathway by Marburg virus VP40: Implications for filovirus biology and budding. PLoS Pathog 16:e1008231.

14. Han Z, Schwoerer MP, Hicks P, Liang J, Ruthel G, Berry CT, Freedman BD, Sagum CA, Bedford MT, Sidhu SS, Sudol M, Harty RN. 2018. Host Protein BAG3 is a Negative Regulator of Lassa VLP Egress. Diseases 6.

15. Liang J, Sagum CA, Bedford MT, Sidhu SS, Sudol M, Han Z, Harty RN. 2017. Chaperone-Mediated Autophagy Protein BAG3 Negatively Regulates Ebola and Marburg VP40-Mediated Egress. PLoS Pathog 13:e1006132.

16. Han Z, Sagum CA, Takizawa F, Ruthel G, Berry CT, Kong J, Sunyer JO, Freedman BD, Bedford MT, Sidhu SS, Sudol M, Harty RN. 2017. Ubiquitin Ligase WWP1 Interacts with Ebola Virus VP40 To Regulate Egress. J Virol 91.

17. Han Z, Sagum CA, Bedford MT, Sidhu SS, Sudol M, Harty RN. 2016. ITCH E3 Ubiquitin Ligase Interacts with Ebola Virus VP40 To Regulate Budding. J Virol 90:9163–71.

18. Shepley-McTaggart A, Fan H, Sudol M, Harty RN. 2020. Viruses go modular. J Biol Chem 295:4604–4616.

19. Klimek C, Kathage B, Wördehoff J, Höhfeld J. 2017. BAG3-mediated proteostasis at a glance. J Cell Sci 130:2781–2788.

20. Anonymous. 2019. WW Domain Proteins in Signaling, Cancer Growth, Neural Diseases, and Metabolic Disorders doi:10.3389/978-2-88963-177-3.

21. Chen YA, Lu CY, Cheng TY, Pan SH, Chen HF, Chang NS. 2019. WW Domain-Containing Proteins YAP and TAZ in the Hippo Pathway as Key Regulators in Stemness Maintenance, Tissue Homeostasis, and Tumorigenesis. Front Oncol 9:60.

22. Lo JY, Chou YT, Lai FJ, Hsu LJ. 2015. Regulation of cell signaling and apoptosis by tumor suppressor WWOX. Exp Biol Med (Maywood) 240:383–91.

23. Abu-Odeh M, Bar-Mag T, Huang H, Kim T, Salah Z, Abdeen SK, Sudol M, Reichmann D, Sidhu S, Kim PM, Aqeilan RI. 2014. Characterizing WW domain interactions of tumor suppressor WWOX reveals its association with multiprotein networks. J Biol Chem 289:8865–80.

24. Rausch V, Hansen CG. 2020. The Hippo Pathway, YAP/TAZ, and the Plasma Membrane. Trends Cell Biol 30:32–48.

25. Dai X, She P, Chi F, Feng Y, Liu H, Jin D, Zhao Y, Guo X, Jiang D, Guan KL, Zhong TP, Zhao B. 2013. Phosphorylation of angiomotin by Lats1/2 kinases inhibits F-actin binding, cell migration, and angiogenesis. J Biol Chem 288:34041–34051.

26. Zhao B, Li L, Lu Q, Wang LH, Liu CY, Lei Q, Guan KL. 2011. Angiomotin is a novel Hippo pathway component that inhibits YAP oncoprotein. Genes Dev 25:51–63.

27. Moleirinho S, Guerrant W, Kissil JL. 2014. The Angiomotins – From discovery to function. 588:2693–2703.

28. Bratt A, Birot O, Sinha I, Veitonmaki N, Aase K, Ernkvist M, Holmgren L. 2005. Angiomotin regulates endothelial cell-cell junctions and cell motility. J Biol Chem 280:34859–69.

29. Moleirinho S, Hoxha S, Mandati V, Curtale G, Troutman S, Ehmer U, Kissil JL. 2017. Regulation of localization and function of the transcriptional co-activator YAP by angiomotin. Elife 6.

30. Ulbricht A, Eppler FJ, Tapia VE, van der Ven PF, Hampe N, Hersch N, Vakeel P, Stadel D, Haas A, Saftig P, Behrends C, Fürst DO, Volkmer R, Hoffmann B, Kolanus W, Höhfeld J. 2013. Cellular mechanotransduction relies on tension-induced and chaperone-assisted autophagy. Curr Biol 23:430–5.

31. Han Z, Ruthel G, Dash S, Berry CT, Freedman BD, Harty RN, Shtanko O. 2020. Angiomotin regulates budding and spread of Ebola virus. J Biol Chem 295:8596–8601.

32. Schuchardt BJ, Mikles DC, Bhat V, McDonald CB, Sudol M, Farooq A. 2015. Allostery mediates ligand binding to WWOX tumor suppressor via a conformational switch. J Mol Recognit 28:220–31.

33. Adler JJ, Johnson DE, Heller BL, Bringman LR, Ranahan WP, Conwell MD, Sun Y, Hudmon A, Wells CD. 2013. Serum deprivation inhibits the transcriptional co-activator YAP and cell growth via phosphorylation of the 130-kDa isoform of Angiomotin by the LATS1/2 protein kinases. Proc Natl Acad Sci U S A 110:17368–73.

34. Adler JJ, Heller BL, Bringman LR, Ranahan WP, Cocklin RR, Goebl MG, Oh M, Lim HS, Ingham RJ, Wells CD. 2013. Amot130 adapts atrophin-1 interacting protein 4 to inhibit yes-associated protein signaling and cell growth. J Biol Chem 288:15181–93.

35. Wang W, Li N, Li X, Tran MK, Han X, Chen J. 2015. Tankyrase Inhibitors Target YAP by Stabilizing Angiomotin Family Proteins. Cell Rep 13:524–532.

36. Choi KS, Choi HJ, Lee JK, Im S, Zhang H, Jeong Y, Park JA, Lee IK, Kim YM, Kwon YG. 2016. The endothelial E3 ligase HECW2 promotes endothelial cell junctions by increasing AMOTL1 protein stability via K63-linked ubiquitination. Cell Signal 28:1642–51.

37. Kim M, Kim M, Park SJ, Lee C, Lim DS. 2016. Role of Angiomotin-like 2 mono-ubiquitination on YAP inhibition. EMBO Rep 17:64–78.

38. Mana-Capelli S, McCollum D. 2018. Angiomotins stimulate LATS kinase autophosphorylation and act as scaffolds that promote Hippo signaling. J Biol Chem 293:18230–18241.

39. Wang C, An J, Zhang P, Xu C, Gao K, Wu D, Wang D, Yu H, Liu JO, Yu L. 2012. The Nedd4-like ubiquitin E3 ligases target angiomotin/p130 to ubiquitin-dependent degradation. Biochem J 444:279–89.

40. Seglen PO, Grinde B, Solheim AE. 1979. Inhibition of the lysosomal pathway of protein degradation in isolated rat hepatocytes by ammonia, methylamine, chloroquine and leupeptin. Eur J Biochem 95:215–25.

41. Dunmore BJ, Drake KM, Upton PD, Toshner MR, Aldred MA, Morrell NW. 2013. The lysosomal inhibitor, chloroquine, increases cell surface BMPR-II levels and restores BMP9 signalling in endothelial cells harbouring BMPR-II mutations. Human Molecular Genetics 22:3667–3679.

42. Licata JM, Simpson-Holley M, Wright NT, Han Z, Paragas J, Harty RN. 2003. Overlapping motifs (PTAP and PPEY) within the Ebola virus VP40 protein function independently as late budding domains: involvement of host proteins TSG101 and VPS-4. J Virol 77:1812–9.

43. Han Z, Madara JJ, Liu Y, Liu W, Ruthel G, Freedman BD, Harty RN. 2015. ALIX Rescues Budding of a Double PTAP/PPEY L-Domain Deletion Mutant of Ebola VP40: A Role for ALIX in Ebola Virus Egress. J Infect Dis 212 Suppl 2:S138–45.

44. Urata S, Noda T, Kawaoka Y, Morikawa S, Yokosawa H, Yasuda J. 2007. Interaction of Tsg101 with Marburg virus VP40 depends on the PPPY motif, but not the PT/SAP motif as in the case of Ebola virus, and Tsg101 plays a critical role in the budding of Marburg virus-like particles induced by VP40, NP, and GP. J Virol 81:4895–9.

45. Ernkvist M, Aase K, Ukomadu C, Wohlschlegel J, Blackman R, Veitonmaki N, Bratt A, Dutta A, Holmgren L. 2006. p130-angiomotin associates to actin and controls endothelial cell shape. FEBS J 273:2000–11.

46. Paramasivam M, Sarkeshik A, Yates JR, 3rd, Fernandes MJ, McCollum D. 2011. Angiomotin family proteins are novel activators of the LATS2 kinase tumor suppressor. Mol Biol Cell 22:3725–33.

47. Chan SW, Lim CJ, Guo F, Tan I, Leung T, Hong W. 2013. Actin-binding and cell proliferation activities of angiomotin family members are regulated by Hippo pathway-mediated phosphorylation. J Biol Chem 288:37296–307.

48. Mana-Capelli S, Paramasivam M, Dutta S, McCollum D. 2014. Angiomotins link F-actin architecture to Hippo pathway signaling. 25:1676–1685.

49. Mercenne G, Alam SL, Arii J, Lalonde MS, Sundquist WI. 2015. Angiomotin functions in HIV-1 assembly and budding. Elife 4.

50. Ray G, Schmitt PT, Schmitt AP. 2019. Angiomotin-Like 1 Links Paramyxovirus M Proteins to NEDD4 Family Ubiquitin Ligases. Viruses 11.

51. Watanabe A, Hippo Y, Taniguchi H, Iwanari H, Yashiro M, Hirakawa K, Kodama T, Aburatani H. 2003. An opposing view on WWOX protein function as a tumor suppressor. Cancer Res 63:8629–33.

52. Pleet ML, Erickson J, DeMarino C, Barclay RA, Cowen M, Lepene B, Liang J, Kuhn JH, Prugar L, Stonier SW, Dye JM, Zhou W, Liotta LA, Aman MJ, Kashanchi F. 2018. Ebola Virus VP40 Modulates Cell Cycle and Biogenesis of Extracellular Vesicles. J Infect Dis 218:S365–S387.

53. Nanbo A, Watanabe S, Halfmann P, Kawaoka Y. 2013. The spatio-temporal distribution dynamics of Ebola virus proteins and RNA in infected cells. Sci Rep 3:1206.

54. Björndal AS, Szekely L, Elgh F. 2003. Ebola virus infection inversely correlates with the overall expression levels of promyelocytic leukaemia (PML) protein in cultured cells. BMC Microbiol 3:6.

55. Wang M, Li Y, Wu M, Wang W, Gong B, Wang Y. 2014. WWOX suppresses cell growth and induces cell apoptosis via inhibition of P38 nuclear translocation in cholangiocarcinoma. Cell Physiol Biochem 34:1711–22.

56. Del Mare S, Salah Z, Aqeilan RI. 2009. WWOX: its genomics, partners, and functions. J Cell Biochem 108:737–45.

57. Salah Z, Aqeilan R, Huebner K. 2010. WWOX gene and gene product: tumor suppression through specific protein interactions. Future Oncol 6:249–59.

58. Chang JY, He RY, Lin HP, Hsu LJ, Lai FJ, Hong Q, Chen SJ, Chang NS. 2010. Signaling from membrane receptors to tumor suppressor WW domain-containing oxidoreductase. Exp Biol Med (Maywood) 235:796–804.

59. Hussain T, Lee J, Abba MC, Chen J, Aldaz CM. 2018. Delineating WWOX Protein Interactome by Tandem Affinity Purification-Mass Spectrometry: Identification of Top Interactors and Key Metabolic Pathways Involved. Front Oncol 8:591.

